# Age at cancer diagnosis by breed, weight, sex, and cancer type in a cohort of over 3,000 dogs: determining the optimal age to initiate cancer screening in canine patients

**DOI:** 10.1101/2022.03.30.486448

**Authors:** Jill M. Rafalko, Kristina M. Kruglyak, Angela L. McCleary-Wheeler, Vidit Goyal, Ashley Phelps-Dunn, Lilian K. Wong, Chelsea D. Warren, Gina Brandstetter, Michelle C. Rosentel, Lisa M. McLennan, Daniel S. Grosu, Jason Chibuk, Dana W.Y. Tsui, Ilya Chorny, Andi Flory

## Abstract

The goal of cancer screening is to detect disease at an early stage when treatment may be more effective. Until recently, cancer screening in dogs has relied upon annual physical examinations and routine laboratory tests, which are largely inadequate for detecting preclinical disease. With the introduction of non-invasive “liquid biopsy” cancer detection methods, the discussion is shifting from “*How* to screen dogs for cancer” to “*When* to screen dogs for cancer”. To address this question, data from 3,452 cancer-diagnosed subjects were analyzed to determine the age at which dogs of certain breeds and weights are typically diagnosed with cancer. In the study population, the median age at cancer diagnosis was 8.8 years, with males diagnosed at younger ages than females, and spayed/neutered dogs diagnosed at significantly later ages than intact dogs. Overall, weight was inversely correlated with age at cancer diagnosis, and purebred dogs were diagnosed at significantly younger ages than mixed-breed dogs. For breeds with 10 or more subjects, a breed-based median age at diagnosis was calculated. A weight-based linear regression model was developed to predict the median age at diagnosis for breeds represented by fewer than 10 subjects and for mixed-breed dogs. The study findings support a general recommendation to start cancer screening for all dogs at the age of 7, and as early as 4 years of age for breeds with a lower median age at cancer diagnosis, in order to increase the chances of early detection and treatment.

## Introduction

Cancer is by far the leading cause of death in adult dogs,^37^ yet options for canine cancer screening have historically been limited in comparison to the robust, guidelines-driven screening programs in human medicine.^109^ With novel cancer screening approaches (such as “liquid biopsy” cancer detection methods using next-generation sequencing of DNA from blood samples) being introduced in veterinary medicine,^25,38,69^ the question of *how* to screen dogs for cancer may soon shift to *when* to start screening dogs for cancer. Currently there is limited evidence to guide veterinarians on this topic, but a “one age fits all” approach to the initiation of screening is unlikely to be appropriate for dogs, given the strong role of both genetic and environmental factors in the development of cancer and the great diversity of breeds and sizes represented in the species.

Previous published studies have often focused on age at cancer diagnosis, or age at death from cancer, for individual breeds ^24,67,127^ or for specific cancer types ^51,61,91^, making the findings difficult to generalize to other breeds or to mixed-breed dogs. Furthermore, some of the larger, population-based studies that incorporated a more diverse selection of breeds were conducted in Europe, ^16,32,47,82^ where “common” breeds may not be representative of breeds that are common in the US; additionally, cancer incidence and cancer types observed in these studies may be different from those seen in a US population, due to environmental differences and spay/neuter rates in Europe versus the US.

The current study examines a large and heterogeneous population of cancer-diagnosed dogs, the vast majority of which were from the US, representing over 120 breeds and a wide variety of cancer types. The purpose of the study is to establish median ages at which dogs of various breeds and weights are diagnosed with cancer, and to develop an evidence-based approach for determining the age at which cancer screening should be initiated for individual dogs.

## Materials and Methods

The study population comprised a total of 3,452 cancer-diagnosed subjects (herein “subjects”) from across three distinct cohorts that contributed data to this study.

The first cohort (“Cohort 1”) comprised subjects prospectively enrolled in the CANcer Detection in Dogs (CANDiD) study.^38^ All subjects were enrolled under protocols that received institutional animal care and use committee (IACUC) or site-specific ethics approval, according to each site’s requirements. All subjects were client-owned, and written informed consent was obtained from all owners.

The subjects in Cohort 1 were all-comers with a current definitive diagnosis of cancer of any type. For these subjects, the dog’s age (known or estimated; in years and months) at enrollment was provided by the enrolling veterinarian, along with a “date of diagnosis” for the patient’s cancer diagnosis. In the case of recurrence of a previous cancer, or a prior history of cancer (before the current diagnosis at enrollment in CANDiD), the “date of diagnosis” represented the date of the first documented cancer diagnosis. Using the dog’s age, date of enrollment, and date of diagnosis, an “age at diagnosis” was calculated for all subjects. Data from a total of 663 subjects from Cohort 1 were used in the current study.

The second cohort (“Cohort 2”) comprised subjects from the National Cancer Institute (NCI) Division of Cancer Treatment and Diagnosis Biological Testing Tumor Repository, deposited by the Canine Comparative Oncology Genetics Consortium (CCOGC).^79^ Samples were prospectively collected from multiple academic institutions within the United States (Colorado State University, The Ohio State University, University of Wisconsin-Madison, Michigan State University, Tufts University, University of California-Davis, University of Missouri, and University of Tennessee), and the Standard Operating Procedures for the collection of samples were approved by each site’s IACUC.

The data from Cohort 2 accompanied clinical samples from dogs with commonly diagnosed cancers, with a focus on enrolling subjects with seven histologies: osteosarcoma, lymphoma, melanoma, hemangiosarcoma, soft tissue sarcoma, mast cell tumor, and pulmonary tumor. Blood samples (collected from subjects prior to surgery) and tumor tissue were collected and submitted to the biorepository. As part of this process, clinical data regarding each subject’s diagnosis were submitted to the NCI. Although the subject’s exact “date of diagnosis” or “age at diagnosis” were not collected as part of this data set, an “age at sample collection” (in years) was documented for each subject. For the purpose of this study, the “age at sample collection” was used as a reasonable approximation for “age at diagnosis”, as the sample collection from the majority of subjects was expected to have occurred within weeks or, at most, months from the time the cancer diagnosis was first made. Data from a total of 1,888 subjects from Cohort 2 were used in the current study.

For Cohorts 1 and 2, breed and weight information were provided for each subject by the veterinarian or staff at the enrolling site.

The final cohort (“Cohort 3”) was a subset of study subjects from a recent publication.^52^ The subjects were patients of the Veterinary Medical Teaching Hospital at the University of California – Davis. Information about each subject was obtained via retrospective chart review and provided in the Supplementary Materials section of the Hart, *et al.* publication; the following data points were used for the purpose of the current study: breed, sex, spay/neuter status, date of birth, date at cancer diagnosis (which allowed for calculation of an “age at diagnosis”), and cancer type (specifically, data was available for four common cancer types: lymphoma, mast cell tumor, osteosarcoma, and hemangiosarcoma). Weight data were not available for any subjects in this cohort. Data from a total of 901 subjects from Cohort 3 were used in the current study.

The overall study population (Cohorts 1, 2, and 3 combined) was examined to determine the mean and median age at cancer diagnosis. Additionally, age at cancer diagnosis was analyzed by breed, weight, sex, and cancer type. P-values were calculated using two-sided t-tests, and p-values of <0.05 were considered statistically significant.

## Results

### Subject demographics

The combined study population of 3,452 subjects comprised 2,537 reported to be purebred, 858 reported to be mixed-breed, and 57 whose breed was described as “other”. As there was no significant difference between the ages at cancer diagnosis for dogs described as mixed-breed and “other” (p=0.6944), these two groups were combined for the purpose of data analysis, resulting in a total of 915 dogs classified as “mixed-breed/other”.

The study population consisted of 1,900 males (55%) and 1,552 females (45%); 76% of males were neutered and 90% of females were spayed. Weight data was available for all 2,551 subjects from Cohorts 1 and 2, and those subjects ranged in weight from 2.5 kg to 98.0 kg, with a mean of 30.3 kg and a median of 30.6 kg (Table 1).

**Table 1:** Demographics of the study population.

| Characteristics | Disposition of study population |
| --- | --- |
| Breed (n=3,452) | Purebred: 2,537 |
|  | Mixed-breed/other: 915 |
| Sex (n=3,452) | Male: 1,900 |
|  | - Male (neutered): 1,452 |
|  | - Male (intact): 446 |
|  | - Male (status not provided): 2 |
|  | Female: 1,552 |
|  | - Female (spayed): 1,390 |
|  | - Female (intact): 161 |
|  | - Female (status not provided): 1 |
| Weight* (n=2,551) | Range: 2.5 – 98.0 kg |
|  | Mean: 30.3 kg |
|  | Median: 30.6 kg |

\* Weight data were not available for subjects in Cohort 3 (n=901)

Subjects were assigned a “cancer type”, based primarily on anatomic location, as previously described.^38^ This classification system was adapted from Withrow and MacEwen’s Small Animal Clinical Oncology (Sixth Edition)^121^ and from the American Joint Committee on Cancer (AJCC) Cancer Staging Manual (Eighth Edition)^11^. The most represented cancer type in the study population was lymphoma, followed by osteosarcoma, mast cell tumor, hemangiosarcoma, and soft tissue sarcoma (Table 2).

**Table 2:** Cancer types* represented in the study population (n=3,452)

| Cancer type | Number of subjects |
| --- | --- |
| Lymphoma/Lymphoid Leukemia | 979 |
| Bone, Osteosarcoma | 664 |
| Mast Cell Tumor | 565 |
| Hemangiosarcoma | 292 |
| Soft Tissue Sarcoma | 240 |
| Malignant Melanoma | 128 |
| Lung | 113 |
| Oral Cavity | 67 |
| Skin | 44 |
| Histiocytic Sarcoma | 40 |
| Peripheral Nerve Sheath | 33 |
| Anal Sac Adenocarcinoma | 29 |
| Multiple Concurrent Primary Cancers | 27 |
| Unknown** | 25 |
| Chondrosarcoma | 22 |
| Liver | 22 |
| Urinary Bladder/Urethra | 18 |
| Nasal Cavity and Paranasal Sinuses | 16 |
| Mammary Gland Carcinoma | 15 |
| Thyroid | 15 |
| Bone, Multilobular Osteochondrosarcoma | 12 |
| Bone, Fibrosarcoma | 9 |
| Adrenal Gland | 8 |
| Bone, Sarcoma (other) | 8 |
| Brain | 8 |
| Spleen | 8 |
| Kidney | 6 |
| Small Intestine | 6 |
| Prostate | 5 |
| Transmissible Venereal Tumor | 5 |
| Heart Base | 3 |
| Pancreas | 3 |
| Bile Duct | 2 |
| Mediastinum | 2 |
| Multiple Myeloma | 2 |
| Salivary Gland | 2 |
| Spinal Cord | 2 |
| Ear Canal | 1 |
| Esophagus | 1 |
| Large Intestine | 1 |
| Nasal Planum | 1 |
| Thymoma | 1 |
| Uterus | 1 |
| Vagina | 1 |

\* Cancer types were classified based primarily on anatomic location, as previously described.<sup>38</sup> \*\* Diagnosis of malignancy was confirmed, but cancer type was not assigned due to limited clinical information.

### Overall distribution of age groups at cancer diagnosis

For the overall cohort of 3,452 dogs, the age at cancer diagnosis ranged from less than one year to 20 years, with a mean of 8.5 years and a median of 8.8 years (Fig. 1).

**Figure 1:**
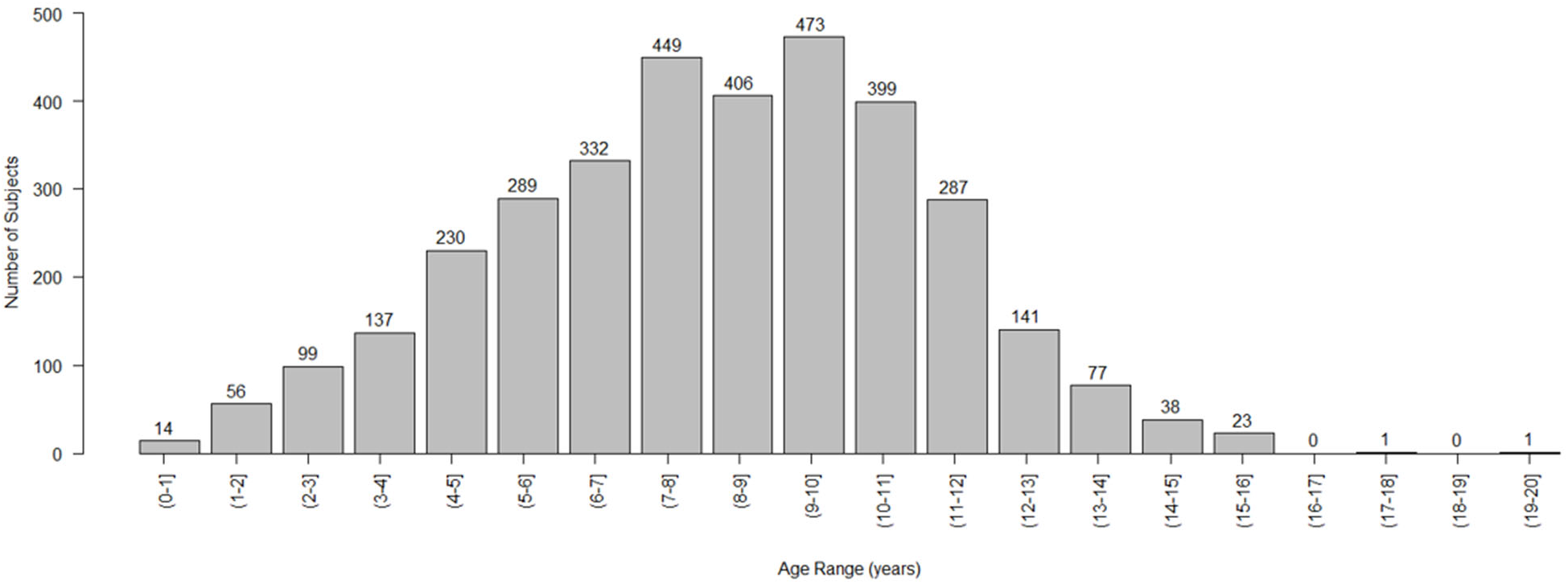
Distribution of subjects by age at cancer diagnosis (n=3,452)

### Age at cancer diagnosis by breed

The age at cancer diagnosis for the 2,537 purebred dogs in this study ranged from less than one year to 20 years of age, with a mean of 8.2 years and a median of 8.0 years. These subjects represented 122 distinct breeds. The most highly represented breeds in the study were Golden Retrievers (n=422) and Labrador Retrievers (n=397), followed by Boxers (n=178), Rottweilers (n=168), and German Shepherds (n=102).

For breeds represented by at least 10 subjects (n=43 breeds), mean and median ages at diagnosis for the breed were calculated. The breeds with the youngest median age at cancer diagnosis were Saint Bernards, Mastiffs, Great Danes, Bulldogs (median: 6 years), followed by Irish Wolfhounds (median 6.1 years), Boxers (median: 6.2 years), and Vizslas and Bernese Mountain Dogs (median: 7.0 years). The breed with the oldest median age at cancer diagnosis was the Bichon Frise (median: 11.5 years) (Fig. 2).

**Figure 2:**
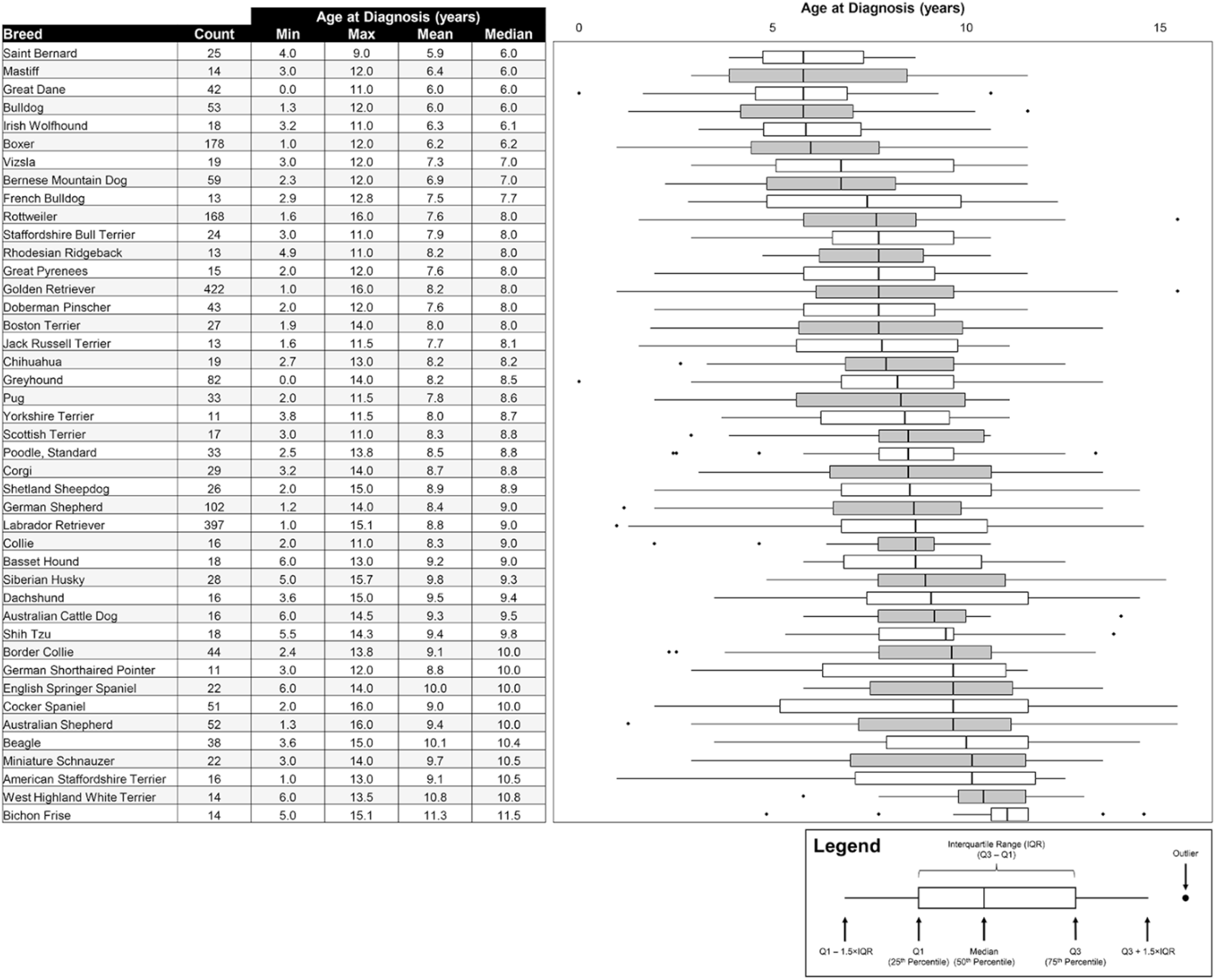
Age distribution at cancer diagnosis by breed (for breeds represented by 10 or more subjects)

For mixed-breed/other dogs (n=915), age at cancer diagnosis ranged from less than one year to 18 years of age, with a mean of 9.2 years and a median of 9.5 years. The mean age at cancer diagnosis for these 915 mixed-breed/other dogs was significantly later than the mean age at diagnosis for the 2,537 purebred dogs in this study (9.2 vs. 8.2 years; p<.0001). (Fig. 3)

**Figure 3:**
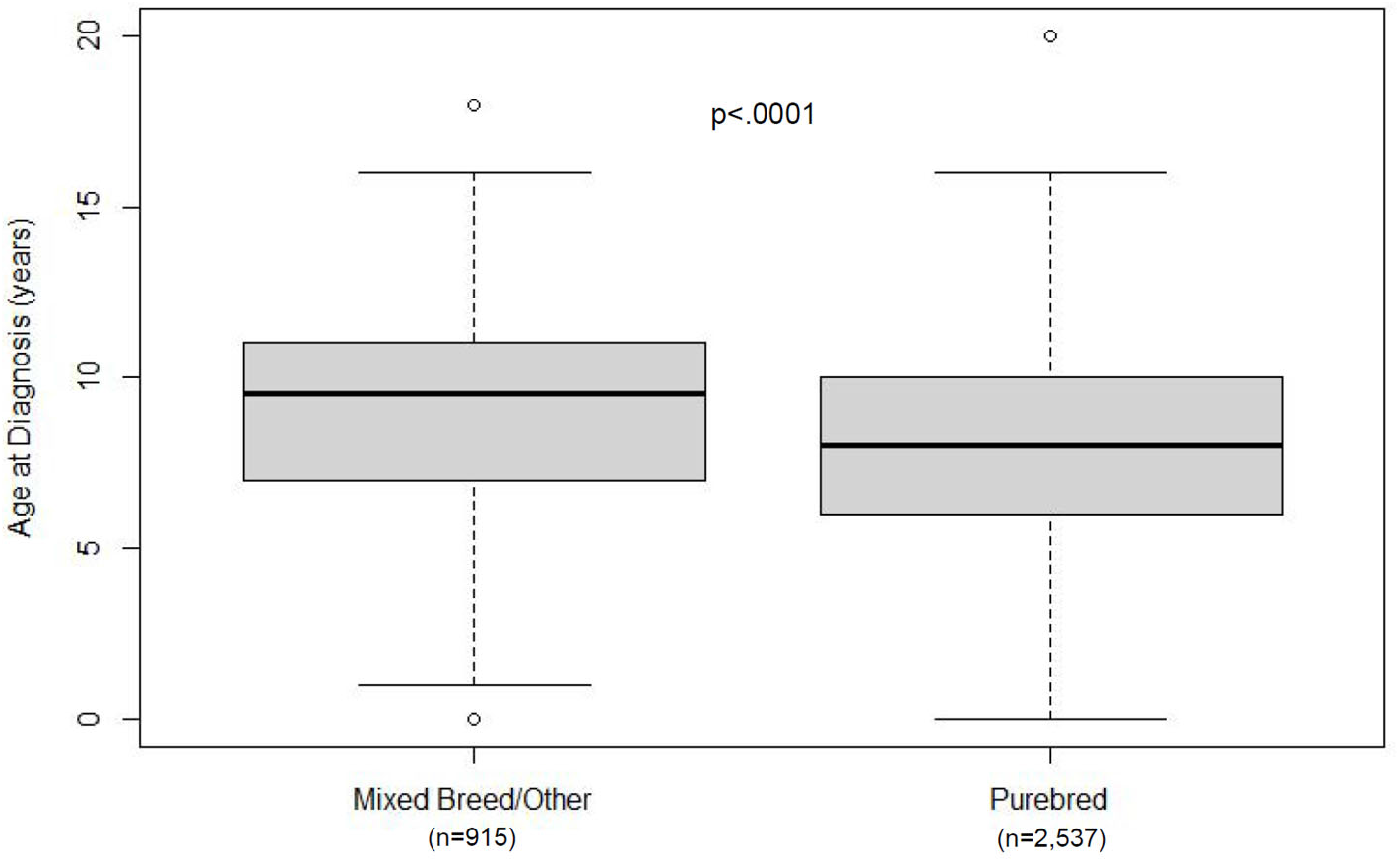
Age distribution at cancer diagnosis for mixed-breed/other vs. purebred subjects.

### Distribution of subjects by weight

Weight data was available for 2,551 subjects. Weight ranged from 2.5 – 98.0 kg (Fig. 4), with a mean of 30.3 kg and a median of 30.6 kg (Table 1).

**Figure 4:**
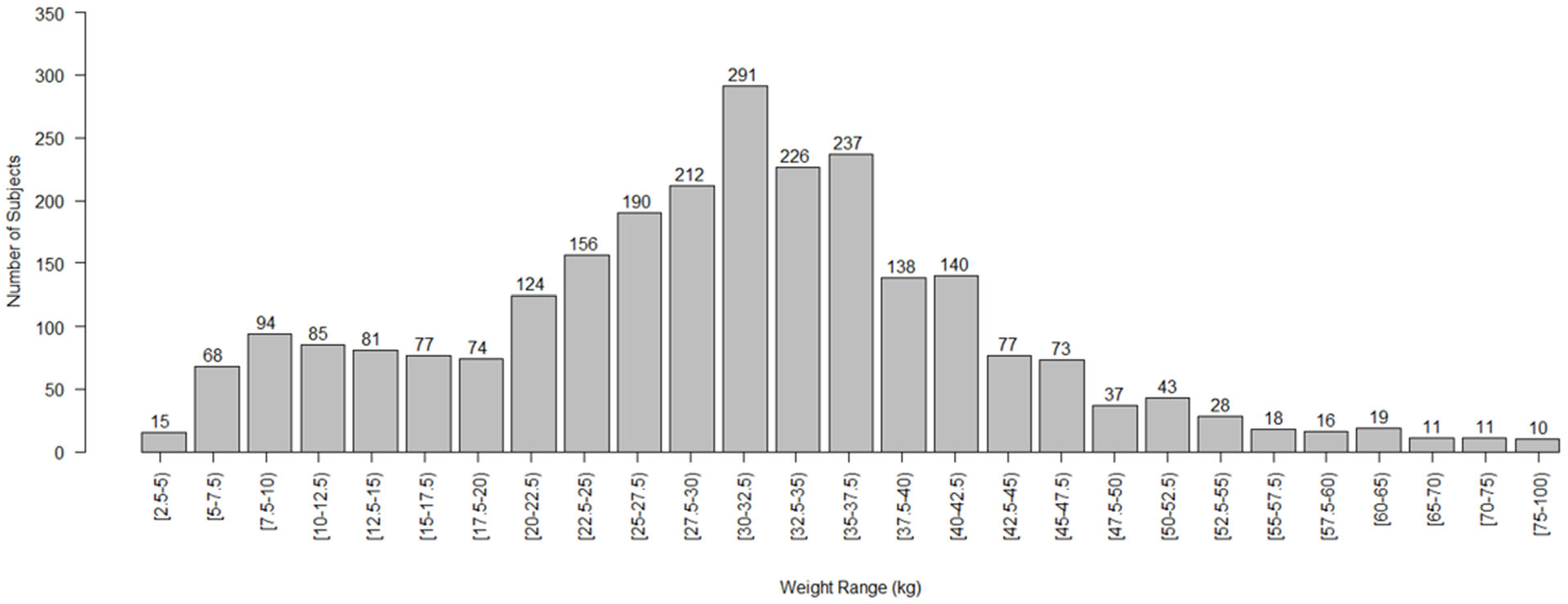
Weight distribution of the study population (for subjects that had a documented weight; n=2,551)

In general, as the weight of the dog increased, the median age at cancer diagnosis decreased. For instance, dogs weighing between 2.5 and 5 kg had a median age at cancer diagnosis of 11 years, compared to 5 years for dogs 75 kg and over. By plotting median age at cancer diagnosis for dogs in various weight brackets, a linear regression formula was derived (herein referred to as the “weight-based model”): Median Age = (−0.068 × Weight) + 11.104 (Fig. 5).

**Figure 5:**
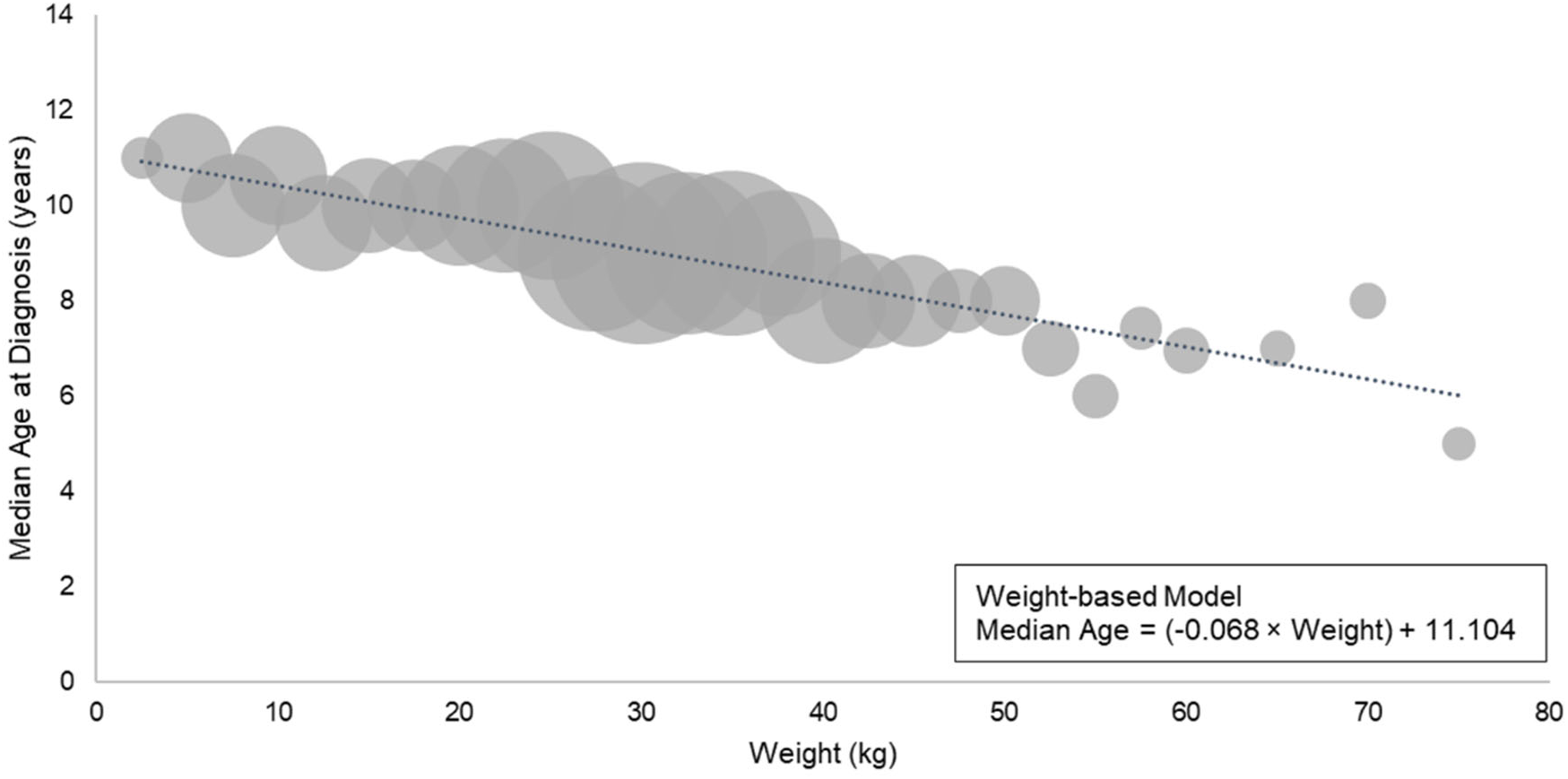
Median age at cancer diagnosis by weight (for subjects that had a documented weight; n=2,551)

Additional analysis was performed on a subset of the breeds from Figure 2. There were 37 breeds for which weight data was available for at least 10 subjects in the study population. For each of these 37 breeds, the median weight of subjects representing that breed in this study was calculated. Then, the actual median age at cancer diagnosis (calculated directly from the subjects of that breed – see Figure 2) was compared to the predicted median age at cancer diagnosis using the weight-based model presented in Figure 5. For the majority of breeds (23 of 37), the median age at cancer diagnosis predicted by the weight-based model was within one year of the actual median age calculated from subjects representing that breed in this study. For certain breeds, particularly Bulldogs, Boxers, Vizslas, French Bulldogs, and Boston Terriers, the median age at cancer diagnosis calculated directly from subjects of that breed was more than 2 years younger than the median age predicted by the weight-based model (Fig. 6).

**Figure 6:**
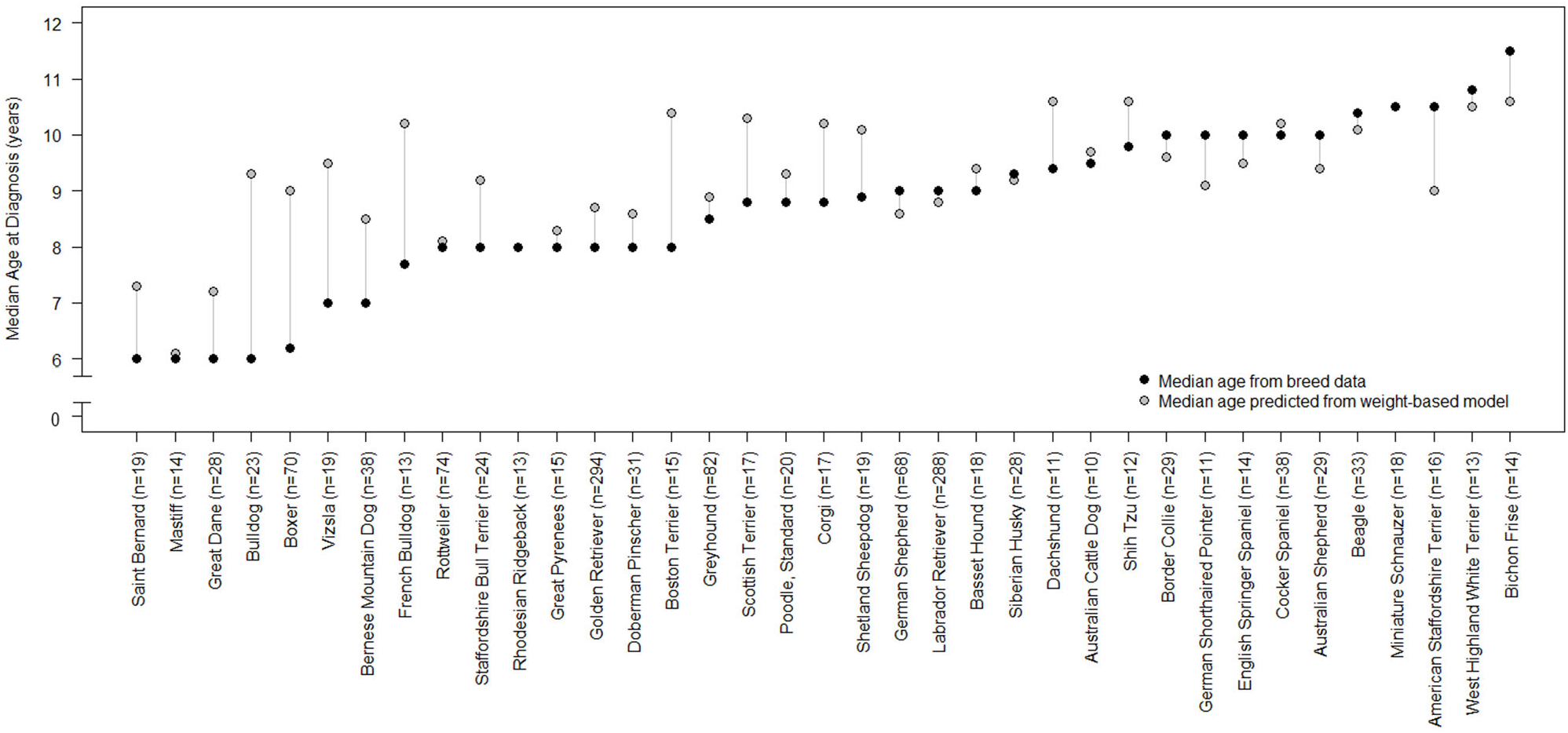
Median age at cancer diagnosis in purebred dogs: breed-based data versus prediction from weight-based model (for breeds that had a documented weight for ≥10 subjects; n=1,495) Please note: Breeds included in Figure 6 are breeds in which weight data were available for 10 or more cancer-diagnosed subjects. Six breeds included in Figure 2 are not represented in Figure 6 due to an insufficient number of subjects with weight information (<10 per breed).

### Age at cancer diagnosis by sex and spay/neuter status

In the overall study population (n=3,452), the age of cancer diagnosis in males was significantly younger than in females (mean 8.3 vs. 8.7 years; p<.0001). When the data were subdivided by sex and spay/neuter status, neutered males were diagnosed with cancer at younger ages than spayed females (mean 8.5 vs. 8.9 years; p=0.0002); however, there was no significant difference between the age at cancer diagnosis for intact males vs. intact females (mean 7.6 vs. 7.3 years; p=0.2623). There was a significant difference between neutered vs. intact males (mean 8.5 vs. 7.6 years; p<0.0001) and spayed vs. intact female dogs (mean 8.9 vs. 7.3 years; p<0.0001), with neutered/spayed dogs showing a significantly later mean age at diagnosis than their intact counterparts (Table 3).

**Table 3:**
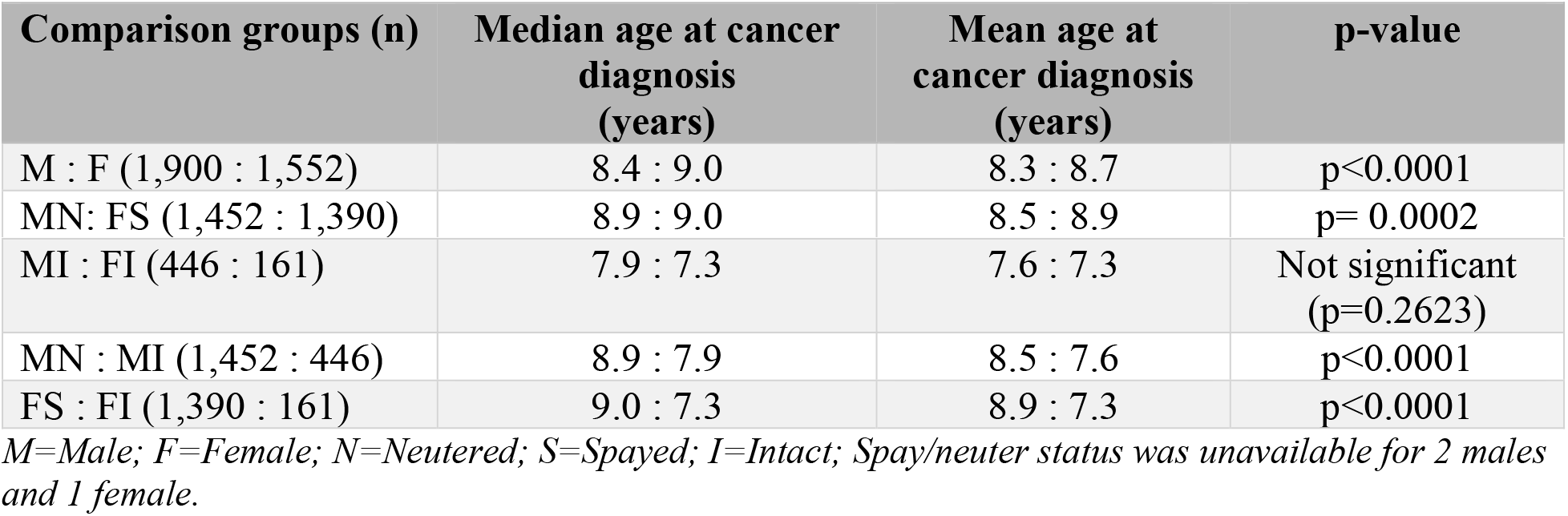
Age at cancer diagnosis by sex and spay/neuter status of the study population.

| Comparison groups (n) | Median age at cancer diagnosis (years) | Mean age at cancer diagnosis (years) | p-value |
| --- | --- | --- | --- |
| M : F (1,900 : 1,552) | 8.4 : 9.0 | 8.3 : 8.7 | $p < 0.0001$ |
| MN: FS (1,452 : 1,390) | 8.9 : 9.0 | 8.5 : 8.9 | $p = 0.0002$ |
| MI : FI (446 : 161) | 7.9 : 7.3 | 7.6 : 7.3 | Not significant<br>(p=0.2623) |
| MN : MI (1,452 : 446) | 8.9 : 7.9 | 8.5 : 7.6 | p<0.0001 |
| FS : FI (1,390 : 161) | 9.0 : 7.3 | 8.9 : 7.3 | p<0.0001 |

M=Male; F=Female; N=Neutered; S=Spayed; I=Intact; Spay/neuter status was unavailable for 2 males and 1 female.

### Age at cancer diagnosis for common cancer types

The median age at cancer diagnosis was analyzed for cancer types represented by at least 10 subjects (n=21 cancer types). Lymphoma/lymphoid leukemia, mast cell tumor, and histiocytic sarcoma all showed median ages at diagnosis younger than age 8; whereas malignant melanoma and cancers of the mammary gland, lung, and urinary bladder/urethra showed median ages at diagnosis of 11 years or older (Fig. 7).

**Figure 7:**
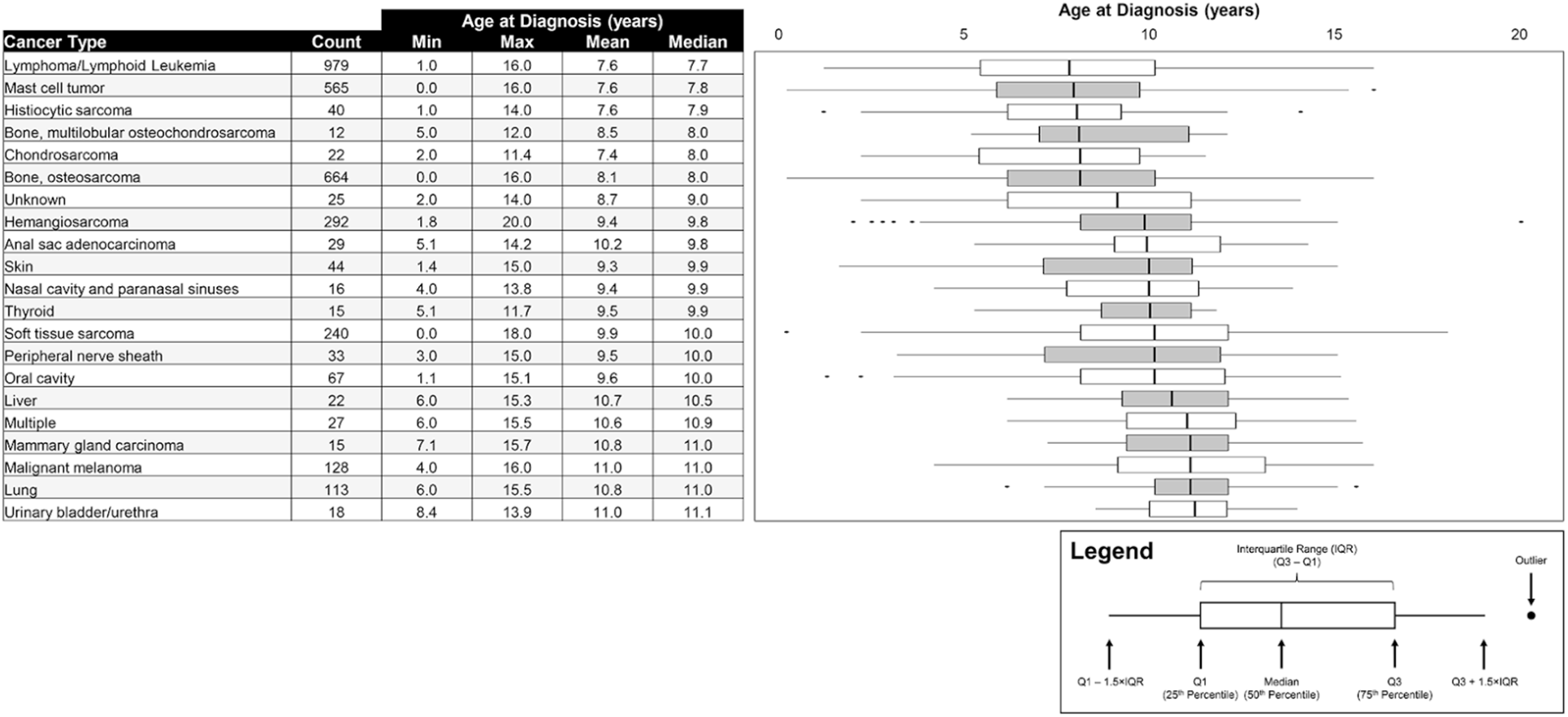
Age distribution at cancer diagnosis by cancer type (for cancer types represented by 10 or more subjects)

## Discussion

Cancer is the leading cause of death in dogs,^37^ and the presence of preclinical malignancy in significant numbers of canine patients has been extensively documented in studies of incidental findings on imaging, surgery, and necropsy.^18,21,27,30,44,99,105,122^ Multiple veterinary professional organizations recognize the value of early cancer detection for optimizing outcomes,^4,9,10,12^ and veterinary academic institutions have issued prevention and screening recommendations for cancer in dogs.^28,29^ However, formal guidelines for earlier detection of cancer through regular screening programs do not currently exist in veterinary medicine as they do in human medicine.^6,38^ It is important to note that the term “screening” is used here in the strict sense,^6,87^ referring to measures taken to detect cancer preclinically in canine patients that are at higher risk for the disease due to age or breed but do not currently have clinical signs indicative of cancer.

Recently, liquid biopsy using next generation sequencing of cell-free DNA was introduced as a novel, non-invasive option for cancer screening in dogs.^38^ With the availability of a blood-based cancer test, the question of *how* to screen dogs for cancer may soon shift to *when* to start screening dogs for cancer.

To address this question, the current study compiled and analyzed data regarding age at cancer diagnosis in dogs of various breeds and weights to help establish evidence-based recommendations for when to initiate cancer screening.

The ages at cancer diagnosis in a population of over 3,400 dogs ranged from <1 year to 20 years, with a median of 8.8 years. Overall, in this study population, males were diagnosed with cancer at younger ages than females, and dogs that had been spayed/neutered were diagnosed at significantly later ages compared to their intact counterparts. The impact of spay/neuter status on the lifetime risk of cancer has been studied previously, with mixed findings. For example, the role of hormonal impact has been well documented in the development of mammary gland carcinoma, with a decreased risk in spayed females compared to reproductively intact females.^106,113^ Conversely, other studies have suggested an increased risk for the development of certain types of cancers in spayed/neutered dogs, such as osteosarcoma, lymphoma, prostate cancer, and bladder cancer.^23,52–54,57,104^ Regardless of the lifetime risk for cancer following spay/neuter, the data from the current study suggest that if cancer is going to develop, it will typically be diagnosed at later ages in dogs that are spayed/neutered.

When age at cancer diagnosis was analyzed by cancer type, the mean and median ages at cancer diagnosis were found to vary significantly across cancer types, with hematological malignancies and mast cell tumors being diagnosed at much younger ages than malignant melanomas and lung cancers. These findings are consistent with previous literature, where the median age at cancer diagnosis for lymphoma has been reported as 6-9 years,^34^ while oral malignant melanoma^101^ and pulmonary tumors^46^ are primarily diseases of older dogs, with reported median ages at diagnosis of 11 years. Furthermore, the mean ages at diagnosis for four common cancers (hemangiosarcoma, lymphoma, mast cell tumor, and osteosarcoma) in the current study are closely aligned with findings from a large study at an academic veterinary center in California, US.^20^

Prior research has found that the lifetime prevalence of common cancers is similar for purebred and mixed-breed dogs, when matched for age, sex, and weight^20^ Building on this research, the current study further evaluated the age at which cancer is diagnosed and found that purebred dogs were diagnosed at significantly younger ages than mixed-breed dogs. This finding could be explained in part by selective breeding methods, which may perpetuate germline mutations that predispose certain breeds to cancer at younger ages.^32^ However, it should be noted that there was wide variability in the age at diagnosis by breed in the purebred cohort in this study, with median ages at diagnosis ranging from 6.0 to 11.5 years (in breeds represented by at least 10 subjects).

In this study, weight was independently correlated with age at cancer diagnosis. Many of the breeds with younger ages at cancer diagnosis in Figure 2 were large- and giant-breed dogs. This finding is supported by Figure 5, which shows that weight appears to be inversely related to age at cancer diagnosis in the overall study population. By plotting a median age at diagnosis for various weight brackets of subjects, a formula for the weight-based model was derived: Median Age = (−0.068 × Weight) + 11.104. This formula can be used to estimate the median age at cancer diagnosis for mixed-breed dogs, or for purebred dogs that have an insufficient number of subjects (<10) to calculate a breed-based median age at diagnosis, if their weight is known.

For the 37 breeds with weight data available from at least 10 subjects, a median weight was calculated for dogs comprising each breed (using demographic data from the study population) and entered into the weight-based model. This allowed the breed-based median age at cancer diagnosis (calculated directly from the subjects of that breed) to be compared to a prediction for median age at cancer diagnosis based only on weight. Though the median age at cancer diagnosis by breed versus weight prediction was similar (within one year) for most breeds, certain breeds showed significant deviations; in particular, breed-based data for Bulldogs, Boxers, Vizslas, French Bulldogs, and Boston Terriers showed median ages at cancer diagnosis at least 2 years younger than the weight-predicted ages. This suggests that genetics may play a stronger role in cancer onset in certain breeds, resulting in younger ages at diagnosis. This aligns with observations from an analysis of cancer claims in more than 1.6 million dogs covered by a leading US pet health insurer over a six-year period, which found significant differences among breeds for both overall relative cancer risk and for average age at first cancer claim. Extensive similarities were noted between the findings of that analysis and those of the present study; for example, Boxers, Great Danes, and French Bulldogs had significantly younger median ages at cancer diagnosis (and average ages at first cancer claim) compared to Beagles, Miniature Schnauzers, and Shih Tzus.^90^

The lifetime risk of cancer as well as cancer mortality in dogs are known to vary significantly by breed.^16,32^ For example, approximately 50% of Irish Water Spaniels and Flat-Coated Retrievers die of cancer, whereas cancer-related mortality is significantly lower in breeds such as Shih Tzus and Dachshunds. However, even in the least-affected breeds, the rate of mortality from cancer is still in the range of 15-20%.^32^ For comparison, common pathophysiologic processes such as traumatic, infectious, metabolic, inflammatory, and degenerative each account for 10% or less of deaths in adult dogs, across all breeds.^37^ Cancer is a leading cause of death even among breeds that are relatively less affected by cancer, suggesting that all dogs – regardless of breed – would derive preventive benefit from regular cancer screening.^38^

Once an approximate age at which cancer may be diagnosed in each dog is calculated, based on breed-specific data or the weight-based model, an “age to initiate screening” can be derived.

Cancer progression timelines are well established in human oncology for multiple types of cancer, and are used to inform recommendations for appropriate screening intervals.^25,88,110,120^ A recent analysis estimated a latency range of 2.2 to 35.7 years for lymphoproliferative and hematopoietic cancers, and 6.6 to 57 years for solid malignancies, with 35 of the 44 cancer types in the analysis found to progress silently for 10 years or longer prior to detection.^85^ Other studies, focusing on genomic evolution timelines across many human cancers, have similarly shown that driver mutations often precede diagnosis by many years to decades.^42^ Analyses focusing on specific cancer types have demonstrated that it takes approximately 17 years for a large benign tumor to evolve into advanced colorectal cancer, but less than 2 additional years for cells within that advanced cancer to acquire metastatic potential;^64^ and in pancreatic cancer, approximately 20 years will pass from the initiation of tumorigenesis until end-stage disease, with metastasis occurring only within the last 2-3 years.^126^ In the recurrence setting, a long-term follow-up study in breast cancer documented recurrence in approximately 25% of patients at distant sites, up to 20 years after the initial curative-intent treatment.^36^

These clinical observations are consistent with tumor growth estimates based on reported tumor doubling times, which have been studied extensively in human cancer. Doubling times of 30 to 300 or more days have been reported for many common cancer types, with significant variation noted across tumor stages, tumor types, and individual patients.^13,33,41,65,80,93,118^ It is generally accepted that a malignant mass becomes clinically detectable (on physical exam or imaging) once it reaches a volume of approximately 1 cm^3^ (1.2 cm diameter), at which point it contains upwards of 1 billion cells and weighs approximately 1 gram.^31,35,45,56^ Using these doubling times, corresponding latency periods can be calculated for cancer in humans, and range from 4 to 25 years, consistent with the clinical studies described above.

Biologically, a similar progression of cancer over an extended period of time is likely to occur in dogs, albeit on a shorter timescale than in humans given the compressed canine lifespan.^25^ Studies of spontaneous and induced canine cancer models have provided estimates of *in vivo* tumor doubling times ranging widely from several days to over 100 days, depending on tumor type and method of measurement, and varying widely across individuals.^19,75,95,96,116^ These doubling times would correspond to latency periods of 1 to 3 years based on the calculation presented above. However, these estimates are likely conservative since cancer is not typically diagnosed as soon as it reaches the threshold of clinical detection; in dogs, cancers are often diagnosed, or present for treatment, in the range of 2.5 to 10 cm^49,62,63,74,78,83,98,105,107,108,117^ (containing 10 billion to 500 billion cells), corresponding to latency periods upwards of 5 years. This estimate is consistent with multi-year latency periods previously documented in dogs following exposure to ionizing radiation: 2 to 10+ years for bone malignancies,^43,59,76^ 2 to 4 years for hemangiosarcomas,^116^ 4 to 10+ years for hepatic malignancies,^48^ and 3 to 10+ years for pulmonary malignancies.^84^

It is also important to note that tumor growth is not linear during the course of cancer progression. Growth tends to be rapid very early in the disease process but slows considerably by the time the disease reaches a clinically detectable size. This progressive increase in the tumor doubling time as the tumor gets bigger is described by Gompertzian growth kinetics (or the Gompertz curve)^71,115^ and is recognized as a feature of both human^1,2,40,50,94^ and canine^14,124^ malignancies. This non-linear growth trajectory further supports the value of general screening, as it implies a relatively long period when the presence of preclinical but detectable cancer could be confirmed by standard clinical evaluation methods, following a positive screening result.

The relatively long duration of cancer progression, in humans and in dogs, affords multiple opportunities for earlier detection over the lifespan through screening at regular intervals.^5,22,35,56,65,80^ In humans, it is recommended to start screening for cancer prior to the peak incidence of age at cancer diagnosis, as noted in breast cancer, where peak incidence occurs in the age group 55-64^86^ and annual or biennial screening mammograms are recommended starting at age 45-50 (or earlier ages for high-risk individuals)^8,39^; or in prostate cancer, where peak incidence occurs in the age group 65-74^86^ and annual or biennial screening is advised to start at age 50 (or earlier ages for high-risk individuals)^7^. Large-scale longitudinal studies are needed to accurately determine the optimal timing and interval of cancer screening in dogs. One such study, the Cancer Lifetime Assessment Screening Study in Canines (CLASSiC) was launched in December 2021 (PetDx, La Jolla, CA); the study aims to prospectively follow over 1,000 initially cancer-free dogs, with semi-annual liquid biopsy testing and comprehensive documentation of cancer-related clinical outcomes, over many years.^26,97^

Considering estimated timeframes for cancer development in dogs, a prudent recommendation would be to start screening for cancer 2 years prior to the median age at diagnosis. In the current study, the median age at diagnosis was close to 9 years (8.8 years), supporting a recommended screening age of 7 years for all dogs. For dogs belonging to breeds with an earlier median age at cancer diagnosis (6 to 7 years), screening should begin as early as 4 years of age. In the current study, 58.3% of subjects (2,012/3,452) were diagnosed with cancer at or before 9 years of age. Indeed, even in breeds with a median age at diagnosis of 10 or greater (Figure 2), 38.0% of subjects (108/284) were diagnosed at or before 9 years of age, reinforcing the benefits of starting to screen no later than age 7 even in those breeds.

This recommendation would align with a screening paradigm centered around a dog’s annual or semiannual wellness visit,^3,5^ with serial testing to increase the opportunity for early detection and intervention. In human cancer screening, the value of repeat (interval) testing is well documented, as it results in a higher cumulative detection rate over the lifespan compared to a single testing event, since each successive test provides an additional opportunity for detection.^66,68,81,128^ A similar scenario is likely to be observed in cancer screening programs for canine patients.^38^

The strengths of this study include the large size of the overall cohort and the wide range of breeds and cancer types represented; however, there are several limitations to acknowledge as well.

For subjects from Cohort 2, the subject’s “age at collection” was used as a proxy for “age at cancer diagnosis”. In doing so, the data presented likely overestimates the actual age at diagnosis, to an unknown extent (possibly by weeks or months). One possible mitigation is that Cohort 2 subjects had their “age at collection” reported in years, rather than years and months, potentially offsetting some of this putative overestimation.

Additionally, the subjects from Cohort 2 and Cohort 3 represented a skewed distribution of cancer types. As noted above, the collection of subjects for Cohort 2 (the Canine Comparative Oncology and Genomics Consortium Biospecimen Repository) was primarily focused on enrolling seven pre-defined cancer types; and Cohort 3 (from the Veterinary Medical Teaching Hospital at the University of California – Davis) has only provided data for four cancer types. These selection biases may have enriched this study for dogs with certain demographic characteristics, because particular cancer types may disproportionately affect dogs of certain breeds, weights or ages; and may have also impacted the estimate for median age at cancer diagnosis for a given breed, if certain cancers were under- or over-represented for that breed in the Cohorts 2 and 3 datasets.

For the cohort of purebred dogs, the median age at cancer diagnosis was calculated for breeds represented by at least 10 subjects. It is unclear whether this is a sufficient number of subjects for deriving a valid median age at cancer diagnosis for each of these breeds. More accurate calculations are expected in the future as larger datasets are collected to inform each of the breed-based estimates.

Another limitation is that breed assignments were provided by the enrolling site, with no way to ensure the accuracy of this information. Furthermore, approximately 2% of dogs were assigned a breed of “other”, with no further information to assign them to a breed category. This small cohort of dogs was combined with the mixed-breed cohort for purposes of data analysis; however, it should be acknowledged that an undefined number of these dogs could be purebred.

Regarding the geographical distribution of subjects, it is estimated that over 95% of dogs in this study were from the United States. This factor may limit the generalizability of the study findings to other geographies in which different environmental characteristics, spay-neuter practices, breed distributions, or other considerations may play a role in age at cancer diagnosis.

Lastly, an assessment of age at diagnosis by breed or weight for specific cancer types was beyond the scope of the current study. This remains an opportunity for future research.

The clinical benefits of earlier cancer detection have been extensively documented in humans^55,60,89,100^ as well as in dogs ^15,17,22,58,70,72,73,77,92,102,103,111,112,114,119,123,125^, and major veterinary professional organizations have emphasized these benefits through statements such as “early detection is critical for the best outcome” and “neoplasia is frequently treatable and early diagnosis will aid [the] veterinarian in delivering the best care possible”.^4,10^ The introduction of novel cancer screening tools for dogs raises the important question of when to start screening for cancer.^38^ This study aims to provide an evidence-based foundation for answering this question by examining the ages at which dogs of various breeds and weights are typically diagnosed with cancer. The study findings support a general recommendation to start screening all dogs at the age of 7, and as early as 4 years of age for breeds that have a lower median age at cancer diagnosis, in order to increase the chances of early detection and treatment. As additional epidemiological data from larger cohorts become available and are incorporated into these algorithms, recommended screening ages can be more accurately determined, particularly for breeds that are underrepresented in this study.

## Acknowledgments

The authors would like to thank all of the dogs (and the humans who love and care for them) enrolled in the various studies that contributed data for this manuscript. Dominique Lau for her assistance creating tables, figures, and graphics to support the visual representation of data associated with this manuscript. Dr. Benjamin L. Hart, Dr. Lynette A. Hart, Abigail P. Thigpen and Dr. Neil H. Willits for the collection and publication of data in the Hart *et al.* 2020 study that were incorporated into the present study as “Cohort 3” (citation 24).

